# A data repository and analysis framework for spontaneous neural activity recordings in developing retina

**DOI:** 10.1101/000455

**Authors:** Stephen John Eglen, Michael Weeks, Mark Jessop, Jennifer Simonotto, Tom Jackson, Evelyne Sernagor

## Abstract

**Background:** During early development, neural circuits fire spontaneously, generating activity episodes with complex spatiotemporal patterns. Recordings of spontaneous activity have been made in many parts of the nervous system over the last 25 years, reporting developmental changes in activity patterns and the effects of various genetic perturbations.

**Results:** We present a curated repository of multielectrode array recordings of spontaneous activity in developing mouse and ferret retina. The data have been annotated with minimal metadata and converted into HDF5. This paper describes the structure of the data, along with examples of reproducible research using these data files. We also demonstrate how these data can be analysed in the CARMEN workflow system. This article is written as a literate programming document; all programs and data described here are freely available.

**Conclusions:** 1. We hope this repository will lead to novel analysis of spontaneous activity recorded in different laboratories. 2. We encourage published data to be added to the repository. 3. This repository serves as an example of how multielectrode array recordings can be stored for long-term reuse.

## Dedication

We dedicate this paper to the memory of our dear friend and colleague Professor Colin Ingram who died December 15th 2013. Colin was the lead investigator on the CARMEN project, from which this study arose.

## Background

The retina is the neural circuit within the eye responsible for converting light signals into neural activity. During at least the first postnatal week of life in the mouse, retinal ganglion cells (RGCs) are spontaneously active, generating waves of activity that propagate across the retina. These spontaneous activity patterns are thought to help refine the development of neuronal connections, as blocking or perturbing the activity leads to altered connectivity patterns. For reviews on the nature and role of spontaneous activity in the nervous system, see [1, 2].

Retinal waves can be studied both using imaging methods and with multielectrode arrays (MEAs). We have collected and annotated these recordings to allow researchers to compare the spatiotemporal properties of recordings obtained from different research groups. We have focused on curating recordings collected by MEAs, as although there are several types of array recording platforms available on the market, the underlying data after spike detection and sorting is simply a set of event times denoting when an action potential was detected on a particular electrode.

This data paper describes the repository we have curated from many key papers investigating the nature of retinal spontaneous activity. We have converted these data files into a common format so that it can be easily shared with others, and provide some scripts to analyse these recordings. We created this repository for several reasons:

1. By building a repository from many sources we are able to effectively compare findings from laboratories acquired under different experimental conditions, and from different transgenic mice.
2. There are few public datasets of MEA recordings, although some are available accompanying research papers [3].
3. We hope this platform will encourage future researchers to contribute their data. Many funding agencies now require data to be archived/shared for several years, and we hope this will serve as an example of how to share data.
4. Converting data to a standard open format, such as HDF5, should ensure that they can be read for many years to come. By contrast, keeping old datasets in proprietary formats may mean that the data are effectively unreadable in a few years.
5. We have used these data as a demonstration for the workflow system in the CARMEN virtual laboratory.

This article is written as an example of “reproducible research”, in that the results should be reproducible by others in a straightforward manner, given the same software and data [4]. The notion of reproducible research is beginning to be practised quite widely in some areas, such as Computational Biology [5], but is not yet that common within most fields, including Neuroscience [6, 7]. Figures and tables marked (Dynamic) in the legends of this article are regenerated dynamically, involving recomputation as needed. (Figures and tables marked (Static) are those that required no computation.) The source file for this article contains LATEX and R code, from which the paper is generated (see section “Availability of supporting data”). Links to all files required to regenerate the paper are provided on the accompanying website [8].

## Data Description

The project web page [8] contains links to the data and code, and will list any updates to the repository. The data are freely available on the CARMEN portal [9]. Free registration to the CARMEN system is required to access the data.

The data provided to us from different laboratories arrived in several text and binary formats. We converted them to one common format to promote their reuse. We chose the open format HDF5 [10] because it provides an efficient and portable framework for storing large datasets. It is supported by many popular computational environments, such as R, Python, Mathematica, Matlab and Julia, and is freely available on all major operating systems. HDF5 is used across many scientific disciplines and is well-tested.

HDF5 stores objects in a hierarchical tree that can be fully specified by the user. Our approach is to store the principal data items (spike times and electrode positions) for a recording in the top level of the tree. Relevant metadata about recordings (such as the age of the retina, and the species) are stored in objects under the /meta/ group of the tree.

### Data format for storage of MEA recordings

The following objects are stored in the root of the HDF5 data files:

1. epos: an *N ×* 2 matrix, where *N* is the number of spike trains in the recording. Row *i* stores the (*x*, *y*) location assigned to spike train *i* in this recording. The values *x* and *y* are specified in µm.
2. sCount: a vector of length *N*. *sCount*[*i*] stores the number of spikes included in spike train *i*.
3. spikes: a vector of length S, where *S* = *sum*(*sCount*). These are the spike times (in seconds). The spike trains for each electrode are concatenated into one long vector, so that the spikes for electrode *j* ∈ [1, *N*] are stored in elements *a* to *b*, where 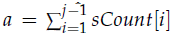 and *b* = *a* + *sCount*[*j*] *−* 1. Within each spike train, the spike times are sorted, smallest first.
4. array: a string describing the MEA used to record the activity. Table 7 lists the values used to date.
5. names: an optional vector of strings of length *N*; *names*[*i*] stores the name assigned to spike train *i*.

#### Summary

To help summarize each recording, we have also created a /summary group containing information which can be readily computed from the spike trains. These summary points can be read from HDF5 files on their own (rather than reading the entire file) and so provides an efficient cache of this information. The following fields are provided.

1. /summary/N: the number of spike trains.
2. /summary/duration: the duration of the recording in seconds, rounded up to the nearest second.
3. /summary/frate: a vector of length *N*. Element *i* stores the firing rate in Hz of spike train *i*.
4. /summary/totalspikes: the total number of spike trains in the file.

## Data sources

Table 2 lists the main studies included in the repository, and the number of files in each collection. A key challenge in creating the repository was writing functions to parse the various formats of source data from the different research groups. This has now been done for each of the major formats. When each data set was converted, tests were performed to check that our results matched those presented in the original publications; some of these checks are discussed later. First we describe each of the key studies included in the repository:

### Blankenship2011

This study investigated the impact of knocking out two connexin isoforms (Cx36 or Cx45) upon spontaneous activity [11]. Recordings of wild-type, single or double (Cx36 and Cx45) knockouts were performed at postnatal day 11/12 (P11/12).

### Demas2003

Recordings over an extended period (P9–P42) were made in wild-type mice, along with mice reared in the dark [12]. This study also served as a baseline for subsequent recordings in transgenic mice [13].

### Demas2006

This set contains data recorded from *nob* mutant mouse, where retinal waves persist at late developmental stages [13].

### Hennig2011

A set of eight recordings investigating the effects of chronic bicuculline application at two critical ages when the effect of GABA upon RGCs switches from excitatory to inhibitory [14].

### Kirkby2013

Recent recordings from wild-type and β2 KO mice recorded in the first postnatal week [15].

### Maccione2014

Wild-type recordings in developing mouse retina recorded from P2 to P13 using a high density (4096 electrode) MEA [16]. This study presents recordings with high spatial resolution obtained at pan retinal scale. It reports that waves become localized hotspots shortly before eye opening and that cellular recruitment within waves increases significantly during the second postnatal week.

### Stacy2005

A set of further control recordings in wild-type mice are provided [17]. In this study, retinal waves were also recorded in transgenic mice where cholinergic neurotransmission was inhibited in most of the retina. However, recordings from the transgenic animal could not be found post-publication.

### Stafford2009

A detailed set of recordings at one age (P6) showing that β2 KO mice do still generate correlated waves [18]; see also [19]. A directional bias in control waves was also reported for the first time.

### Sun2008

Two different versions of the β2 KO transgenic mice were studied, and in comparison to earlier studies [20], shown to have correlated activity extending over larger distances than wild type. Although the key paper [19] focuses on postnatal days 4 and 5, recordings from a range of days are provided, and were analysed separately [21].

### Torborg2004

This key summarises data that appeared in several publications [20, 22–24]. These were the first MEA recordings of spontaneous activity in the β2 KO mouse, and showed that although individual RGCs were spontaneously active, the correlations in firing of neighbouring RGCs were strongly reduced. There are also recordings combining the β2 KO line with a gap junction knockout (CX36), and recordings under different pharmacological conditions.

### Wong1993

These are the first MEA recordings of retinal waves, in ferret at different developmental ages. (Some data from here were presented also in [25].) This was the first paper to introduce the key measure of correlated activity, the correlation index, that has been subsequently used in most studies. Although the recordings are relatively short, they highlighted strong distance-dependent correlations gradually decaying with age. (Conversion of these files was complicated as they were binary files, so we converted them through a custom macintosh program written for these data by Markus Meister.)

### Xu2011

To investigate further the effect of cholinergic neurotransmission, a transgenic line (where β2-nAChR was expressed in just RGCs of β2 KO mice) was generated [26]. In this β2(TG) line, waves were restored, although the spatial extent of correlations was reduced.

### Citing the data

We are grateful to our colleagues for sharing their recordings. If you use any of these data sets, we request that you acknowledge the relevant authors by citing the corresponding papers (listed in Table 2).

### Minimal metadata

Our approach to metadata is deliberately minimal, simply describing what we think are some of the essential features of the recordings, such as developmental age, and genotype. This of course means that many details are missing, but in most cases we hope that they can be extracted (manually) from the corresponding publications. We store the metadata as named items in the HDF5 file, under the /meta/ group. For example, the developmental age of the recording is stored in /meta/age. Table 1 describes the metadata that are included in the HDF5 files.

**Table 1.**
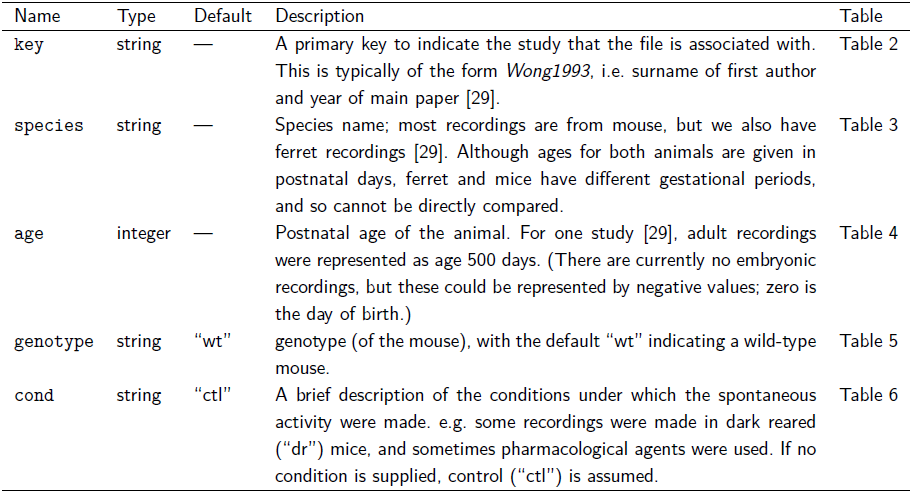
Metadata items stored in the repository. Values with no default are compulsory. The final column refers to subsequent tables for more details on each name. (Static)

| Name | Type | Default | Description | Table |
| --- | --- | --- | --- | --- |
| key | string | — | A primary key to indicate the study that the file is associated with. This is typically of the form *Wong1993*, i.e. surname of first author and year of main paper [29]. | Table 2 |
| species | string | — | Species name; most recordings are from mouse, but we also have ferret recordings [29]. Although ages for both animals are given in postnatal days, ferret and mice have different gestational periods, and so cannot be directly compared. | Table 3 |
| age | integer | — | Postnatal age of the animal. For one study [29], adult recordings were represented as age 500 days. (There are currently no embryonic recordings, but these could be represented by negative values; zero is the day of birth.) | Table 4 |
| genotype | string | “wt” | genotype (of the mouse), with the default “wt” indicating a wild-type mouse. | Table 5 |
| cond | string | “ctl” | A brief description of the conditions under which the spontaneous activity were made. e.g. some recordings were made in dark reared (“dr”) mice, and sometimes pharmacological agents were used. If no condition is supplied, control (“ctl”) is assumed. | Table 6 |

**Table 2.** Keys and citations of the data sources in the repository. n is the number of files associated with each key. (Dynamic)

| key | n | citation |
| --- | --- | --- |
| Blankenship2011 | 17 | [11] |
| Demas2003 | 37 | [12] |
| Demas2006 | 41 | [13] |
| Hennig2011 | 8 | [14] |
| Kirkby2013 | 22 | [15] |
| Maccione2014 | 48 | [16] |
| Stacy2005 | 7 | [17] |
| Stafford2009 | 29 | [18] |
| Sun2008 | 62 | [19] |
| Torborg2004 | 54 | [22] |
| Wong1993 | 11 | [29] |
| Xu2011 | 30 | [26] |

**Table 3.** Numbers of recordings for each species. (Dynamic)

| species | n |
| --- | --- |
| ferret | 11 |
| mouse | 355 |

**Table 4.** Numbers of files from each study (rows) for each postnatal age (columns). (Dynamic)

|  | 0 | 1 | 2 | 3 | 4 | 5 | 6 | 7 | 8 | 9 | 10 | 11 | 12 | 13 | 15 | 21 | 30 | 42 | 500 |
| --- | --- | --- | --- | --- | --- | --- | --- | --- | --- | --- | --- | --- | --- | --- | --- | --- | --- | --- | --- |
| Blankenship2011 | 0 | 0 | 0 | 0 | 0 | 0 | 0 | 0 | 0 | 0 | 0 | 17 | 0 | 0 | 0 | 0 | 0 | 0 | 0 |
| Demas2003 | 0 | 0 | 0 | 0 | 0 | 0 | 0 | 0 | 0 | 2 | 0 | 5 | 0 | 3 | 9 | 8 | 0 | 10 | 0 |
| Demas2006 | 0 | 0 | 0 | 0 | 0 | 0 | 0 | 10 | 0 | 0 | 0 | 0 | 10 | 0 | 8 | 13 | 0 | 0 | 0 |
| Hennig2011 | 0 | 0 | 0 | 0 | 0 | 2 | 2 | 0 | 0 | 2 | 2 | 0 | 0 | 0 | 0 | 0 | 0 | 0 | 0 |
| Kirkby2013 | 0 | 0 | 0 | 0 | 2 | 7 | 10 | 3 | 0 | 0 | 0 | 0 | 0 | 0 | 0 | 0 | 0 | 0 | 0 |
| Maccione2014 | 0 | 0 | 1 | 2 | 4 | 4 | 6 | 2 | 4 | 7 | 5 | 7 | 3 | 3 | 0 | 0 | 0 | 0 | 0 |
| Stacy2005 | 0 | 0 | 0 | 0 | 0 | 7 | 0 | 0 | 0 | 0 | 0 | 0 | 0 | 0 | 0 | 0 | 0 | 0 | 0 |
| Stafford2009 | 0 | 0 | 0 | 0 | 0 | 0 | 29 | 0 | 0 | 0 | 0 | 0 | 0 | 0 | 0 | 0 | 0 | 0 | 0 |
| Sun2008 | 0 | 1 | 3 | 2 | 13 | 17 | 3 | 4 | 4 | 4 | 2 | 6 | 1 | 2 | 0 | 0 | 0 | 0 | 0 |
| Torborg2004 | 0 | 0 | 0 | 2 | 12 | 9 | 0 | 1 | 3 | 0 | 16 | 11 | 0 | 0 | 0 | 0 | 0 | 0 | 0 |
| Wong1993 | 1 | 1 | 0 | 0 | 1 | 1 | 0 | 0 | 0 | 0 | 0 | 0 | 1 | 0 | 2 | 2 | 1 | 0 | 1 |
| Xu2011 | 0 | 0 | 0 | 0 | 30 | 0 | 0 | 0 | 0 | 0 | 0 | 0 | 0 | 0 | 0 | 0 | 0 | 0 | 0 |

**Table 5.** Numbers of recordings of each genotype included in the repository. Note that Wong1993 data are from ferret. (Dynamic)

|  | Beta2 KO | Beta2 KO/Cx36 KO | Beta2 Picciotto | Beta2 Xu | Beta2(TG) | Cx36 KO | Cx36 KO/Cx45 KO | Cx45 KO | nob | wt |
| --- | --- | --- | --- | --- | --- | --- | --- | --- | --- | --- |
| Blankenship2011 | 0 | 0 | 0 | 0 | 0 | 2 | 6 | 4 | 0 | 5 |
| Demas2003 | 0 | 0 | 0 | 0 | 0 | 0 | 0 | 0 | 0 | 37 |
| Demas2006 | 0 | 0 | 0 | 0 | 0 | 0 | 0 | 0 | 25 | 16 |
| Hennig2011 | 0 | 0 | 0 | 0 | 0 | 0 | 0 | 0 | 0 | 8 |
| Kirkby2013 | 11 | 0 | 0 | 0 | 0 | 0 | 0 | 0 | 0 | 11 |
| Maccione2014 | 0 | 0 | 0 | 0 | 0 | 0 | 0 | 0 | 0 | 48 |
| Stacy2005 | 0 | 0 | 0 | 0 | 0 | 0 | 0 | 0 | 0 | 7 |
| Stafford2009 | 0 | 0 | 0 | 0 | 0 | 0 | 0 | 0 | 0 | 29 |
| Sun2008 | 0 | 0 | 21 | 18 | 0 | 0 | 0 | 0 | 0 | 23 |
| Torborg2004 | 11 | 7 | 0 | 0 | 0 | 19 | 0 | 0 | 0 | 17 |
| Wong1993 | 0 | 0 | 0 | 0 | 0 | 0 | 0 | 0 | 0 | 11 |
| Xu2011 | 0 | 0 | 0 | 0 | 17 | 0 | 0 | 0 | 0 | 13 |

**Table 6.** Pharmacological and environmental conditions under which spontaneous activity were recorded. Where possible, the descriptions of the conditions follow those provided with the original data. (Dynamic)

| cond | n |
| --- | --- |
| 10 microM forskolin | 1 |
| 2 microM forskolin | 2 |
| 2 microM strychnine | 1 |
| 2.88 microM FPL | 2 |
| 25 microM APV + 25 microM CNQX;<br>wash then 50 microM Mecamylamine | 1 |
| 25 microM APV + 50 microM CNQX | 1 |
| 50 microM Carbenoxolone | 1 |
| bic | 4 |
| ctl | 332 |
| dr | 19 |
| wash | 2 |

**Table 7.** Descriptions of the MEA layouts in the repository, along with their name and number (n) of recordings. (Dynamic)

| array | n | description |
| --- | --- | --- |
| MCS_8x8_100um | 154 | 60 electrodes in square lattice (corners missing) with 100 μm spacing (Multi Channel Systems). |
| MCS_8x8_200um | 70 | 60 electrodes in square lattice (corners missing) with 200 μm spacing (Multi Channel Systems). |
| APS_64x64_42um | 48 | 4096 electrodes in square lattice with 42 μm spacing (L Berdondini, IIT). |
| litke_hex_60um | 29 | 512 electrodes on hexagonal lattice with 60 μm spacing (AM Litke, UCSC). |
| EJC1_hex_60um | 54 | 61 electrodes on hexagonal lattice with 60 μm spacing (EJ Chichilnisky, Salk). |
| stanford_hex_60um | 11 | 61 electrodes on hexagonal lattice with 60 μm spacing (Stanford). |

**Table 8.** Summary of spike sorting methods used to create spike trains. (Static)

| Key | Spike sorting method |
| --- | --- |
| Blankenship2011 | Plexon offline sorter |
| Demas2003 | Plexon offline sorter |
| Demas2006 | Plexon offline sorter |
| Hennig2011 | Wave_clus [43] |
| Kirkby2013 | Plexon offline sorter |
| Maccione2014 | None |
| Stacy2005 | None |
| Stafford2009 | Mixture of Gaussians model [44] |
| Sun2008 | Plexon Offline sorter |
| Torborg2004 | Manual clustering |
| Wong1993 | Manual clustering |
| Xu2011 | Plexon offline sorter |

#### Sorting

We attempted to include metadata regarding spike detection and sorting. However, many different methods for spike detection and sorting have been used in the last 25 years. These vary from manual, semi-automated to fully automated. Furthermore, the level of details included in publications varied significantly. Rather than encode these in the metadata, we instead provide a brief textual summary of the methods in Table 8.

## Analyses

We now provide some examples of reproducible research using the repository. The aim is to summarise the main features of the repository, rather than to provide novel analyses of these data.

### R package

We have used the R programming environment to develop a package of tools for the analysis of spontaneous activity. R, however, is not required to use these data files. This R package (called sjemea) was created in 2001 (to support work subsequently published [12]) and is still under development. The package primarily focuses on the batch analysis of data; other open tools more suitable towards interactive analysis are available [27, 28]. We now give several examples of analysing the repository using code from this R package.

### Overview of the repository

Figure 1 provides an overview of the repository, showing that recordings range from just a few minutes to many hours. In particular, we see that two different groups recorded waves continuously for up to eight hours [14, 17].

**Figure 1.**
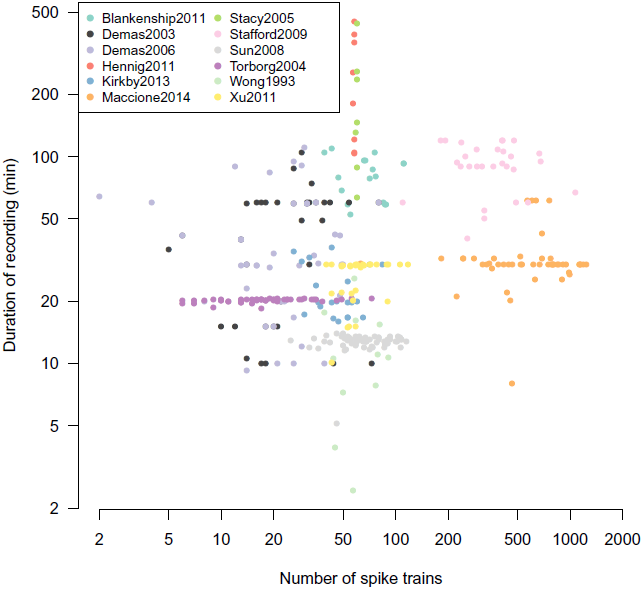
Basic features of recordings in the repository. Each recording is summarised by its number of spike trains and its duration. The colour of each point indicates which collection the recording comes from. Both axes are plotted on a log scale. We currently have 366 recordings in the repository, occupying 298 MB on disc. (Dynamic)

#### Units

As all recordings were from extracellular electrodes, each electrode can detect activity from multiple neurons. Most, but not all, recordings have been spikesorted to discriminate activity from multiple on each electrode. We therefore refer to spike trains coming from “units” throughout this paper to avoid confusion between multi-unit activity from several neurons and inferred activity from a single neuron. Most recordings contain fewer than 100 spike trains because they were made on MEAs consisting of 60 or 64 electrodes. There are also two clear groups of recordings with around 500–1300 units which were recorded from the two higher-density arrays [16, 18].

### Fourplot

The “fourplot” is our one page summary plot of a recording which we use as an initial screen to check its quality. Figure 2 shows one such example. This plot allows us to quickly evaluate the recording using the features described in the figure legend. The accompanying website has a gallery section showing the fourplot for each datafile in the repository.

**Figure 2.**
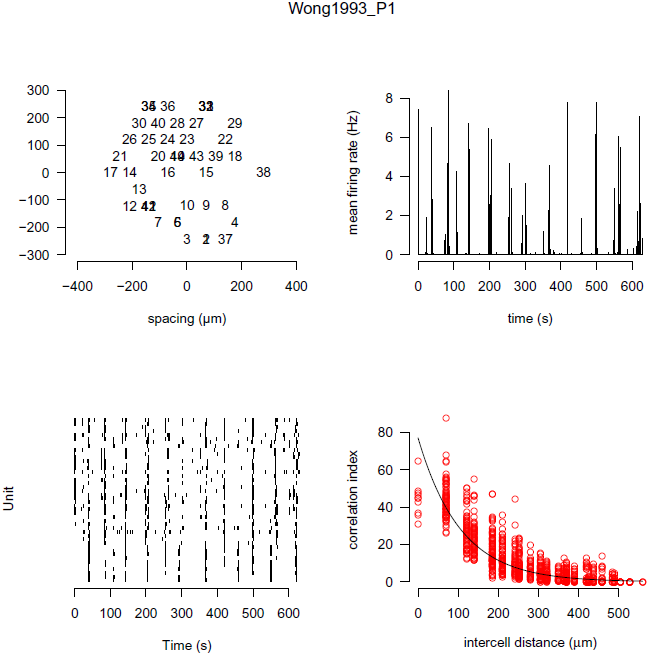
Ten minutes of spontaneous activity from P1 ferret retina recorded using a MEA. The name of the data file is given at the top of the plot. Top left: the estimated position of each unit is plotted. Each spike train is given a unique number; overlapping numbers indicate that more than one unit was assigned the same position. Top right: the firing rate estimated in one second bins, averaged across the entire array. Periodic elevations in firing rate, followed by long periods of relative silence, are characteristic of retinal waves. Bottom left: the raster showing the spike times of all units (starting with unit one at the bottom). Bottom right: the correlation index plot [29], described in detail in Figure 3. (Dynamic)

### Correlation indices in neonatal ferret retina

Retinal waves induce correlations in the firing patterns of neighbouring RGCs. This was first demonstrated in the analysis of ferret retinal waves using the correlation index measure [29]. The correlation index measures the degree that two neurons spike together within some small time window (typically 50 ms). Figure 3 shows the correlation index as a function of the distance separating any given pair of electrodes. This figure almost exactly replicates Figure 8 of the prior study [29], with the only exception that we also show correlation indices for pairs of neurons with zero distance separating them.

**Figure 3.**
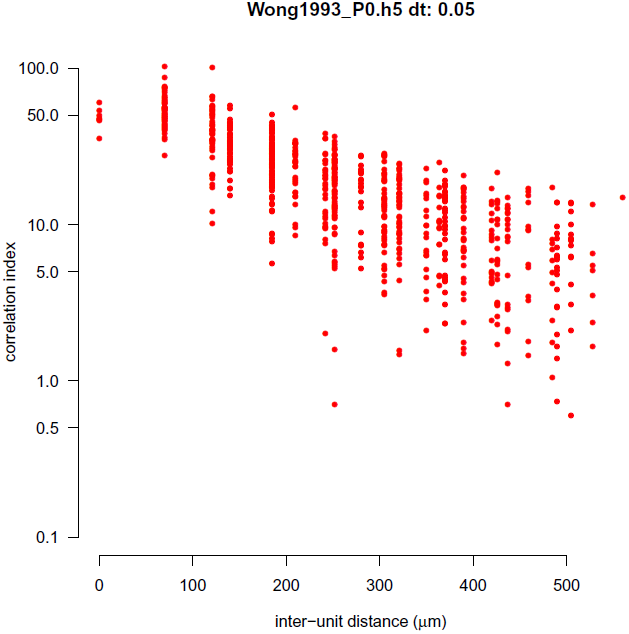
Example correlation index plot. For each pair of units we plot the correlation index computed between the two spike trains against the distance on separating the two units. This plot denotes the correlation index curve for the file Wong1993_P0.h5 (ferret P0), and matches the correlation index plot shown in Figure 8 of [29]. (The y axis is plotted on a logarithmic axis to match the original figure.) The only observable difference is that in our plot we have included correlations for pairs of units that share the same array location (inter-unit distance = 0 µm). (Dynamic)

### Correlation indices in wild-type and transgenic mice

Cholinergic neurotransmission is required for the generation of retinal waves in early development [30, 31]. One key transgenic line has been global knockout of the β2 subunit of the nicotonic acetylcholine receptor (nAChR), termed β2 KO here. Initial reports suggested that β2 KO mice lack retinal waves [20, 32], but subsequent studies reported retinal waves in these mice [18, 19]. These differences might have occurred because of different recording conditions, notably temperature and bath medium [18]. Given the importance of these results, we have collected and curated most of the key recordings published to date that quantify spontaneous activity in β2 KO mice.

The effects of transgenically rescuing β2-nAChR into just RGCs, leaving it knocked out in the rest of the nervous system, were recently investigated [26]. In this genetic manipulation, termed β2(TG), retinal waves were correlated over shorter distances than waves in wild-type mice (Figure 1G of [26]). We have recreated that result by recalculating the correlation indices. Figure 4 has the same key properties as previously reported with the notable exception that the correlation indices in both wild-type and β2(TG) mice are about half the magnitude compared to those originally reported. The shape of the two groups, and the overall conclusions, are unaffected however. The discrepancy between the two results is likely to be an artifact of the method previously used [26], although their code is no longer available to confirm this.

**Figure 4.**
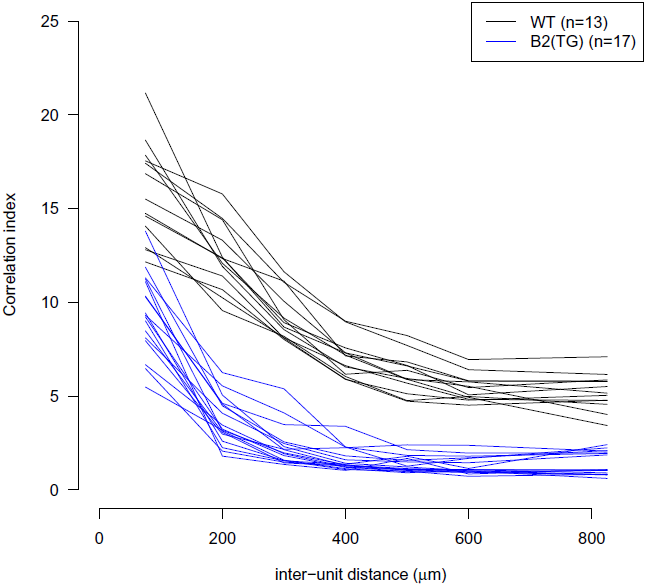
Family of correlation index plots to summarise retinal wave data recorded at P4 [26]. Each line represents the mean correlation index, where the inter-unit distance has been grouped into approximately 100 µm bins, from one recording. The colour of the line denotes whether the recording was taken from a wild-type or β2(TG) mouse. (Dynamic)

### CARMEN application: burst analysis

We have made our data freely available in the CARMEN system [33]. The CARMEN Virtual Laboratory is a collaborative online facility for neuroscientists. Data can be uploaded to a repository, and shared with other neuroscientists. Extensive metadata [34] can be attached to the data, and a search facility allows data to be located in the repository. The system is currently targeted towards electrophysiology data, and predominantly MEA and EEG data.

Useful neuroscience analysis routines can be converted into CARMEN services [35]. Service code can be written in a range of programming languages (including Matlab, Python, R, C/C++, Java), and can be easily wrapped into a service using the CARMEN Service Builder tool. Metadata attached to each service provides information for the user and the system. Once a service has been registered with the CARMEN system, users can run them via the portal, using any available data on the repository. The service is executed within the CARMEN system’s execution environment, which is a private cloud of heterogeneous servers. The execution environment can support multiple execution server environments; currently these are Windows Server, Centos5 Linux and Scientific Linux 4 platforms. The execution details are hidden from the user.

The CARMEN portal also deploys a workflow tool within the browser, to allow users to tie services together into a processing pipeline. To support the interaction of services, a common data standard for all data types is used; the Neural Data Translation Format (NDF). NDF provides a standard format for neural time series data, segment data, and event data [36] and has a rich metadata header which provides a detailed description of the data contents, and which supports the addition of annotations and other relevant attachments (e.g. visual or audio files). A workflow editor provides a graphical means to construct and edit workflows. A workflow enactment engine allows the workflow to be run over the execution servers. The heterogeneous nature of the service infrastructure means that a workflow can be constructed from services that are written using different programming languages and for differing platforms.

To demonstrate the virtual laboratory workflows on these data, three services were built in Matlab and compiled into a standalone executable for inclusion into the service framework:

1. HDF5 to NDF converter — This reads in the HDF5 file and converts it into an NDF neural event data file.
2. A burst detection service — This finds bursts independently within each spike train of a recording [14]. The service takes input data in the form of an NDF neural event file.
3. A graphing service to plot burst durations of multiple input files.

The CARMEN workflow facility (Figure 5) chains these services together so that given an input file, it is first converted into NDF and then the burst times are computed. The output from the independent burst analysis services are then compared to generate a plot such as Figure 6. This figure demonstrates that median burst duration is around 0.1 s and there is good agreement between recordings from different laboratories, albeit with one recording showing a few bursts longer than 2 s.

**Figure 5.**
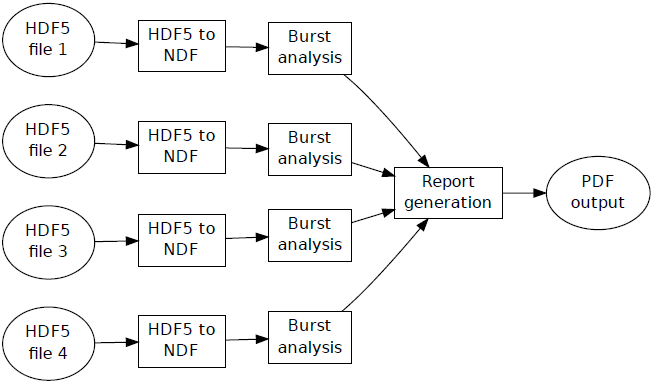
Example of the CARMEN workflow facility. Ellipses denote files and boxes denote CARMEN services. Four HDF5 files are independently converted into NDF for burst analysis. The outputs (in this case, the duration of each burst detected on each electrode) are then collated into a report generation service to produce a graphical summary, shown in Figure 6. (Static)

**Figure 6.**
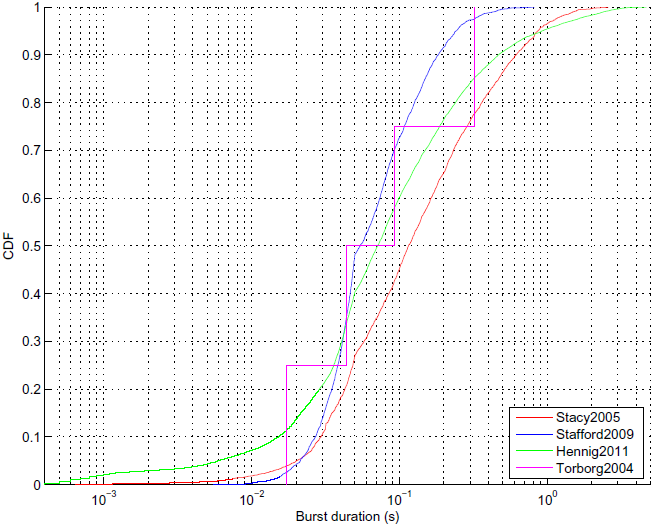
Burst durations for spontaneous activity recorded from four laboratories at postnatal day 5 or 6. This file is the output from the CARMEN workflow illustrated in Figure 5. One file was selected from four different datasets. Each curve shows the cumulative distribution (denoted CDF) of burst durations. The number of bursts detected from each recording were Stacy2005: 2903 bursts (from 60 units); Stafford2009: 3338 (571); Hennig2011: 16896 (58); Torborg2004: 4 (13). (Static)

## Discussion

### Role of the repository

Given the ongoing debate about whether neuronal activity instructs the development of neuronal circuits [37, 38], we believe it is important to understand the spatiotemporal properties of waves in different recordings. This repository provides a framework for systematic studies of spontaneous activity, looking for example at the effects of particular mutations, such as the β2 KO. Furthermore, we can now begin to study the variability between laboratories when recording spontaneous activity in the retina, as has already been reported for cortical cultures [39]. We hope this repository will also lead to increased data reuse.

### Collecting new data

We hope that the repository will prove useful and we encourage investigators to provide recordings of spontaneous activity. We typically require just spike times (rather than voltage traces) and a description of how to map neurons to positions on the array. The minimal metadata is also required, typically in a spreadsheet. It is preferable if the data have already been presented in an article, so that we can refer to that article. Unpublished data can also be accepted as long as the investigator is aware that the data are made freely available.

The current focus of the repository is on collecting spontaneous activity from developing retina. Since spontaneous activity is present in other systems, we also anticipate extending the repository to include data from e.g. cortical and hippocampal cultures [40].

### Generating new standards

To date, there are no established standards for storing spike trains recorded from MEAs; there are a range of data formats from different hardware vendors which our R package can process. However, one key aim of the datasharing program of the International Neuroinformatics Coordinating Facility (INCF) is to establish such standards. We hope that the data provided here can be used as a useful test case for evaluating any proposed standards [41]. Given the relatively simple format in which our data are currently provided, we imagine that changing the data files to accommodate any new standards should be straightforward.

## Methods

All data reported in this paper have been previously published, see Table 2. Details of the experimental procedures are available in those articles. None of the recordings have previously been made freely available. In all cases, recordings of spike times (rather than voltage traces) were provided. Files were then converted into the common HDF5 framework described in this paper. During this conversion, data were checked where possible with previous reports, e.g. descriptions of mean firing rates. Metadata were validated using a separate script. The fourplot (Figure 2) for each recording was also checked. In a few instances this led to discussions with the original authors, or some data being excluded.

## Availability of supporting data

The HDF5 files are available as a zip file [9]; accompanying code is linked to from the project web page [8]. This article is an example of a literate programming document. It has been created in R using the knitr package [42]. All the figures and tables in this paper are generated dynamically as the document is compiled. Several R packages are required to run the analysis. Full details are given on the “Code” section of the accompanying website.

## List of abbreviations used

CARMEN: Code Analysis, Repository and Modelling for E-Neuroscience
CDF: Cumulative Distribution Function
CRAN: Comprehensive R Archive Network
EEG: Electroencephalography
HDF5: Hierarchical Data Format, version 5
INCF: International Neuroinformatics Coordinating Facility
MEA: multielectrode array
NDF: Neural Data Translation Format
Pn: Postnatal day *n*, e.g. P5 for Postnatal day 5
RGC: retinal ganglion cell

## Authors’ contributions

- Conceived and designed the project: SJE, ES.
- Collected and pre-processed data from groups: SJE, JDS.
- Curated and processed data: SJE.
- Contributed code: SJE.
- CARMEN workflow analysis: MJ, TJ, MW.
- Wrote the manuscript: SJE, TJ, ES, MW.

All authors read and approved the final manuscript.

## Acknowledgements

We thank all the investigators who have contributed data to the repository. Thanks to Matthias Hennig and Matthew Down for providing a burst analysis algorithm for use in the CARMEN workflow. Andrew Morton provided a Neuroexplorer script for converting data files. This work was supported by grants from EPSRC (EP/E002331/1), BBSRC (BB/H023577/1 and BB/I000984/1) and Wellcome Trust (083205/B/07/Z).

## Session information

- R version 3.0.2 (2013-09-25), `x86_64-apple-darwin10.8.0`
- Locale: `en_GB.UTF-8/en_GB.UTF-8/en_GB.UTF-8/C/en_GB.UTF-8/en_GB.UTF-8`
- Base packages: base, datasets, graphics, grDevices, methods, parallel, stats, utils
- Other packages: knitr 1.5, RColorBrewer 1.0-5, rhdf5 2.6.0, sjemea 0.40, xtable 1.7-1
- Loaded via a namespace (and not attached): digest 0.6.4, evaluate 0.5.1, formatR 0.10, highr 0.3, stringr 0.6.2, tools 3.0.2, zlibbioc 1.8.0

Paper Revision: 1.62

